# Retinotopic biases in contextual feedback signals to V1 for object and scene processing

**DOI:** 10.1101/2024.03.26.586553

**Authors:** Matthew A. Bennett, Lucy S. Petro, Clement Abbatecola, Lars Muckli

## Abstract

Identifying the objects embedded in natural scenes relies on recurrent processing between lower and higher visual areas. How is cortical feedback information related to objects and scenes organised in lower visual areas? The spatial organisation of cortical feedback converging in early visual cortex during object and scene processing could be retinotopically specific as it is coded in V1, or object centred as coded in higher areas, or both. Here, we characterise object and scene-related feedback information to V1. Participants identified foreground objects or background scenes in images with occluded central and peripheral subsections, allowing us to isolate feedback activity to foveal and peripheral regions of V1. Using fMRI and multivoxel pattern classification, we found that feedback of object information is projected to foveal V1 cortex with increased detail during an object identification task. Background scene information is projected to both foveal and peripheral V1 but can be disrupted by a sufficiently demanding object discrimination task. We suggest that the feedback connections during scene perception project back to earlier visual areas an automatic sketch of occluded information to the predicted retinotopic location. In the case of a cognitive task however, feedback pathways project content to foveal retinotopic space, potentially for introspection, functioning as a cognitive active blackboard and not necessarily predicting the object’s location. This feedback architecture could reflect the internal mapping in V1 of the brain’s endogenous models of the visual environment that are used to predict perceptual inputs.

## Introduction

The brain acquires knowledge about the outside world by generating predictions of upcoming sensory input, sending predictions in cortical feedback pathways down to lower hierarchical stages and learning from prediction failures (Rao and Ballard, 1999; Lee and Mumford, 2003; Friston, 2005). The cortical visual system is composed of reciprocally connected, hierarchically organised areas. This hierarchical network encodes internal models of the visual world and uses cortical feedback signals to contextualise or predict feedforward information (Pennartz et al., 2019). Following this account, prediction signals, originating in areas that respond with spatial invariance, would restore spatial precision when fed back to sensory areas, because feedforward processing in early visual cortex is organised in a precise retinotopic map of visual space. Therefore, cortical feedback input to sensory areas could reflect a precise, reverse topography of feedforward processing. However, cortical microcircuitry provides evidence that feedback input to sensory areas could also reflect the more abstract form of higher visual areas and be liberated from retinotopic coordinates (Shipp, 2016). For example, Wang et al., (2022) observed a systematic variation of feedback connectivity to V1 and V2 linking central/upper visual field and lower visual field to ventral and dorsal stream areas, respectively, which could represent an interface between higher and lower-level encoding.

Retinotopy at one level of the cortical hierarchy is inherited from the previous level, and in turn reciprocal feedback connections target neurons with similar retinotopic preferences (Salin et al., 1992) to amplify or disamplify the signal locally. One example of this is that figure boundaries are detected in V1 during the feedforward sweep, and subsequent excitatory feedback signals from retinotopically-matched neurons enhance the representation of the figure surface (Roelfsema et al., 2002). Hence, higher areas could use feedback processing to sketch a spatially precise interpretation of the stimulus by leveraging V1’s high spatial resolution capabilities. Another example is border ownership, where topographic feedback projections sent down the cortical hierarchy disambiguate which side of a contour belongs to an object and which side belongs to background (Self et al., 2019). However, one consideration for the hypothesis that feedback targets neurons with reciprocal feedforward connections is that cortical feedback pathways also reach locations in lower areas (i.e., V1 or V2) where no feedforward stimulus is simultaneously processed. For example, novel objects presented in the periphery induce foveal activity (in a distinct location from where objects were presented) relevant for the task of comparing the object properties (Williams et al., 2008). This finding suggests that cortical feedback can also liberate itself from the retinotopic coordinate system to engage early visual cortex in a way more suited to cognitive processing requirements. David Mumford (1991) suggested that the early visual cortex functions as an ‘active blackboard’, where specialised higher areas project back to early areas a sketch of the content of their scene segmentation. This concept of visual processing can be developed to include a cognitive space, with the early visual cortex acting as a cognitive active blackboard (Roelfsema and de Lange, 2016). Here, more specialised areas for higher cognitive processes project the content of mental operations back down to early visual areas, for example, when comparing two novel objects presented in the periphery, the features of these objects are projected to foveal workspace. The results from Williams et al. (2008) inspired a series of studies exploring this peripheral to foveal feedback using TMS and behavioural paradigms (reviewed in Oletto et al., 2022). Similarly, when scene images are partially occluded, specialised areas project back internal model predictions similar to mental line drawings to the occluded region (Morgan et al., 2019). Beyond the visual system, Falchier et al. (2002) found cross-modal auditory projections targeting peripheral but not foveal V1. In a functional MRI study, we found contextual auditory scene information to project to the peripheral retinotopic visual areas but less so to the foveal regions (Vetter et al., 2014; Vetter et al. 2020).

Here, using 3T human fMRI, we investigated the role of primary visual cortex V1 during the discrimination of peripheral objects, during the simultaneous automatic processing of scene information. We tested whether feedback signals related to objects or scenes were directed to foveal versus peripheral V1 and how task influences this processing. In two experiments, we presented images containing objects, background scenes, or a combination of the two a similar approach to Harel et al. (2013), but now, by occluding central and peripheral subsections of the image, we are able to isolate feedback activity in the corresponding foveal and peripheral subsections of V1. This approach allows us to test the retinotopic biases in contextual feedback signals to V1 for object and scene processing. In the first experiment, similar objects as used by Williams et al. (2008) were superimposed onto unrelated background scenes. In the second experiment, objects appeared embedded in congruent naturalistic scenes. In both experiments, participants performed an object or scene discrimination task.

## Results

### Experiment 1

We investigated if information about novel objects presented to the periphery is fed back to foveal V1 (Williams et al., 2008) when objects are overlaid on scenes. We studied feedback information for objects and scenes in V1 by analysing multivoxel information patterns in the cortical representations of occluded subsections of four images. The images were one of two grayscale natural scenes (‘Mountain’ and ‘Seaweed’) with a pair of superimposed abstract objects belonging to one of two object categories (‘Cubic’ and ‘Smooth’, Figure 1). We occluded central and lower right image portions to prevent informative feedforward input to foveal and peripheral subsections of V1. We classified either the scene or object identity in foveal and peripheral non-stimulated subsections of V1, independently for scene and object tasks.

**Figure 1.**
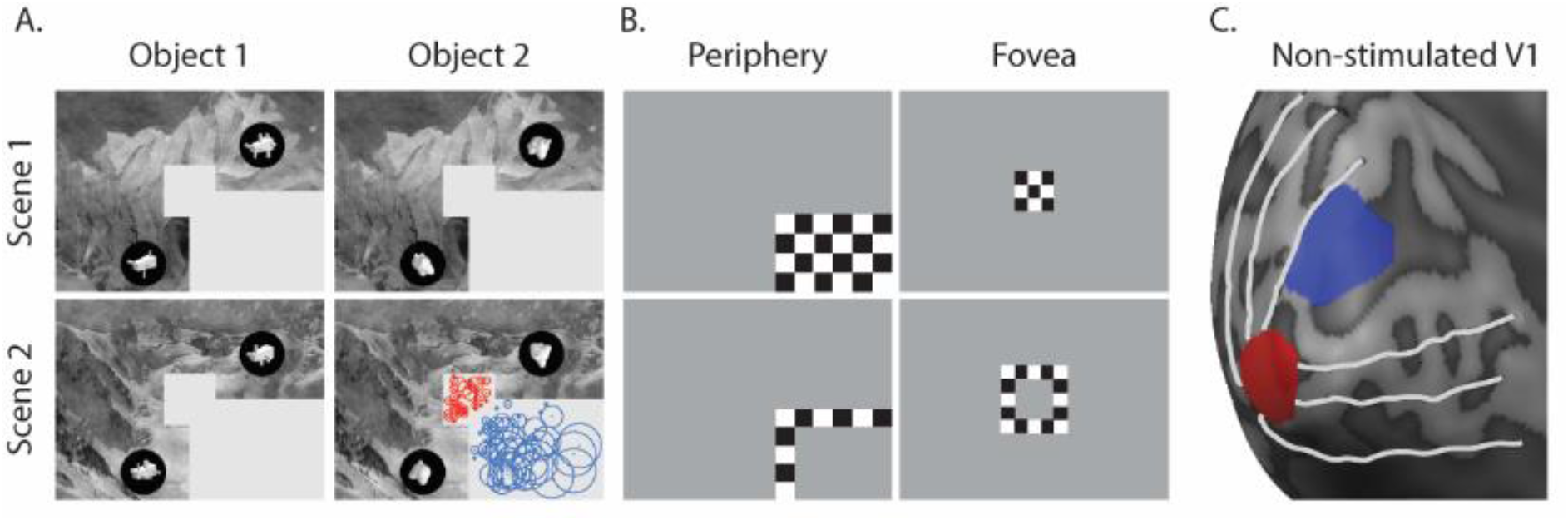
**A**. The experimental images presented in experiment 1, with an example of the estimated receptive fields of the V1-ROI overlaid. **B**. ROI mapping stimuli. For each occluded ROI, we contrasted the checkerboard in the top with the checkerboard in the bottom row to produce an initial ROI (periphery in blue, fovea in red) as shown in C. We then further restricted this ROI to those voxels with pRFs falling entirely within the occluded region (example of pRFs shown in A for object 2 and scene 2).

### Behaviour

Participants performed one of two tasks while fixating centrally; judging whether the two objects were identical or not, or indicating the scene identity when cued by a frame that was blurred. When participants decided if the two peripheral objects were identical or not, they scored 76.6% (± 13.2% stderr) correct. During the background scene task, participants detected the blurred frames 74.4% (± 7.8% stderr) of the time, with a false alarm rate of 25.1% (± 3.8% stderr). The scene task could be completed without compromising global attention to the scene and matched the difficulty of the object task. After detecting the blurred frame, participants were good at identifying the background scene: 92.1% (± 0.8% stderr).

### fMRI brain imaging – MVPA

To investigate the feedback processing in occluded V1 regions we used multivoxel pattern classifiers on BOLD responses from occluded foveal and peripheral voxels. We were able to decode scene information during the scene task only, in both foveal (Scene Task: 54.6%, *p*<0.01; Object Task: 49.6%, *p*=0.658) and peripheral V1 (Scene Task: 54.7% p<0.01; Object Task: 52.5%, *p*=0.099, Figure 2). At the foveal V1 location, we found that correct scene classification rates were significantly lower during the object task than the scene task (p=0.01). This result suggests that scene information is fed back to foveal and peripheral V1, but that the demanding task of perceptually discriminating abstract objects disrupts the representation of scene information.

**Figure 2.**
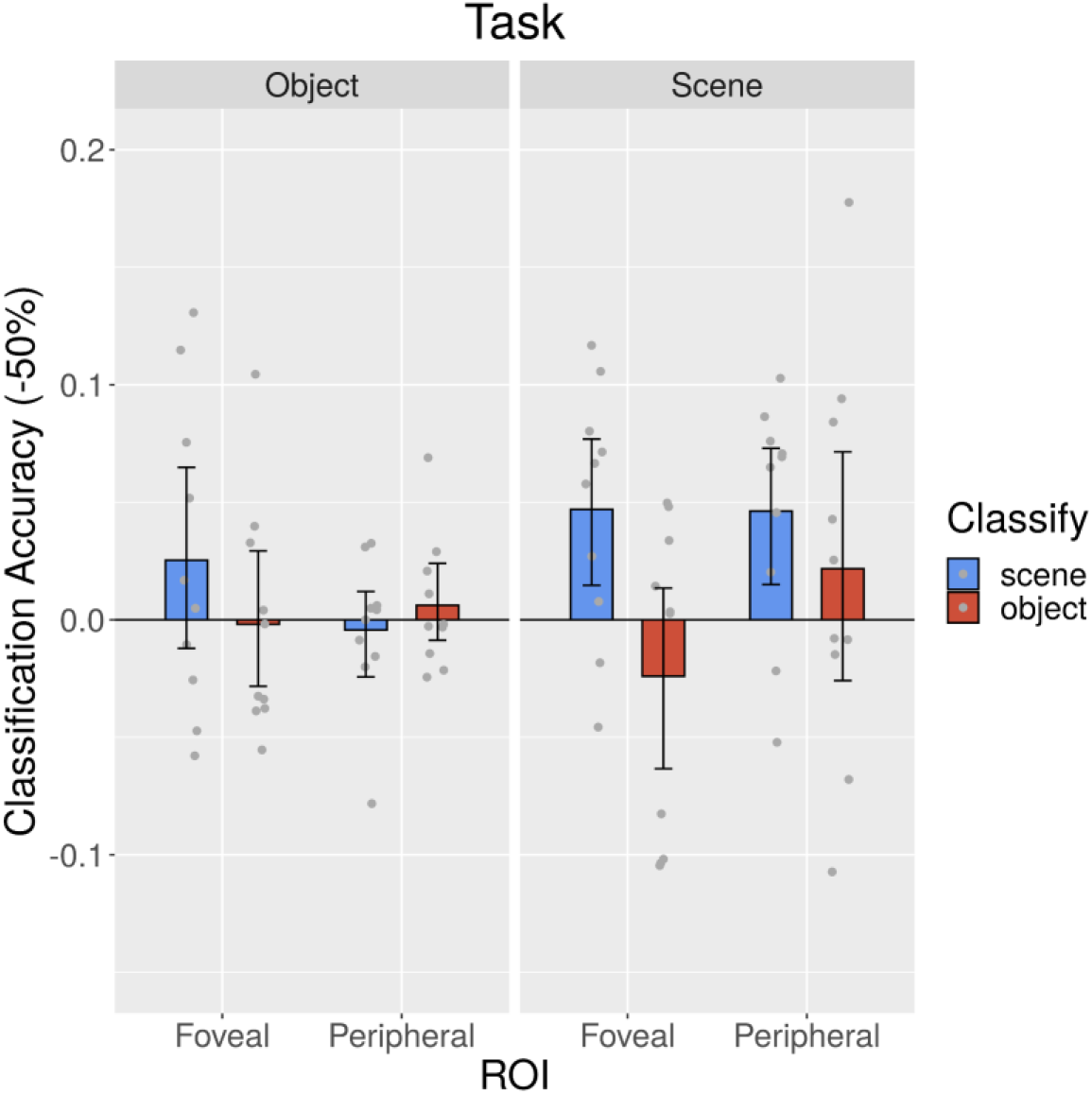
Multivariate pattern analysis (MVPA) classification accuracies for all conditions of experiment 1, in foveal and peripheral regions of interest in V1, during object and scene tasks.

We were not able to detect object identity information in either task in foveal (Object Task: 50.6%, *p=*0.242; Scene Task: 52.2%, *p*=0.194) or peripheral V1 (Object Task: 49.8%, *p*=0.568; Scene Task: 47.6%, *p*=0.888, Figure 2). This contrasts with Williams et al. (2008) who were able to decode object identity information in the fovea. In our data, even classification in a V1 ROI directly stimulated in a feedforward manner by the objects was low and only significantly above chance during the object task (Object Task: 52.1%, p<0.001; Scene Task: 48.6%, p=0.961). This seemingly low feedforward performance could be because we generated unique object instances for every trial. This design means that the pattern classifier had to generalise across object categories rather than rely on particular retinotopic features. Williams et al. used the same strategy and observed a within category correlation during the feedforward condition of r = 0.29 versus r = 0.24 in the feedback condition. Given that our objects were unnatural in appearance and superimposed onto the scene in a superficial way, and given that discriminating the objects disrupted scene feedback, we reasoned that the scene feedback information could have obscured any object feedback information, even in regions processing the objects. To investigate this possibility, we conducted experiment 2 where we used images of objects and scenes with a realistic context.

### Experiment 2

In experiment 2, we used computer generated grayscale images of real objects embedded in scenes in a natural way, for example, a barbeque on a beach (Figure 3). The object task was to identify the object in the image (BBQ, Tent, or None). Similarly, in the scene task participants had to identify the background scene in the image (Beach, Forest or None). We localised peripheral and foveal non-stimulated ROIs in V1 as in experiment 1.

**Figure 3.**
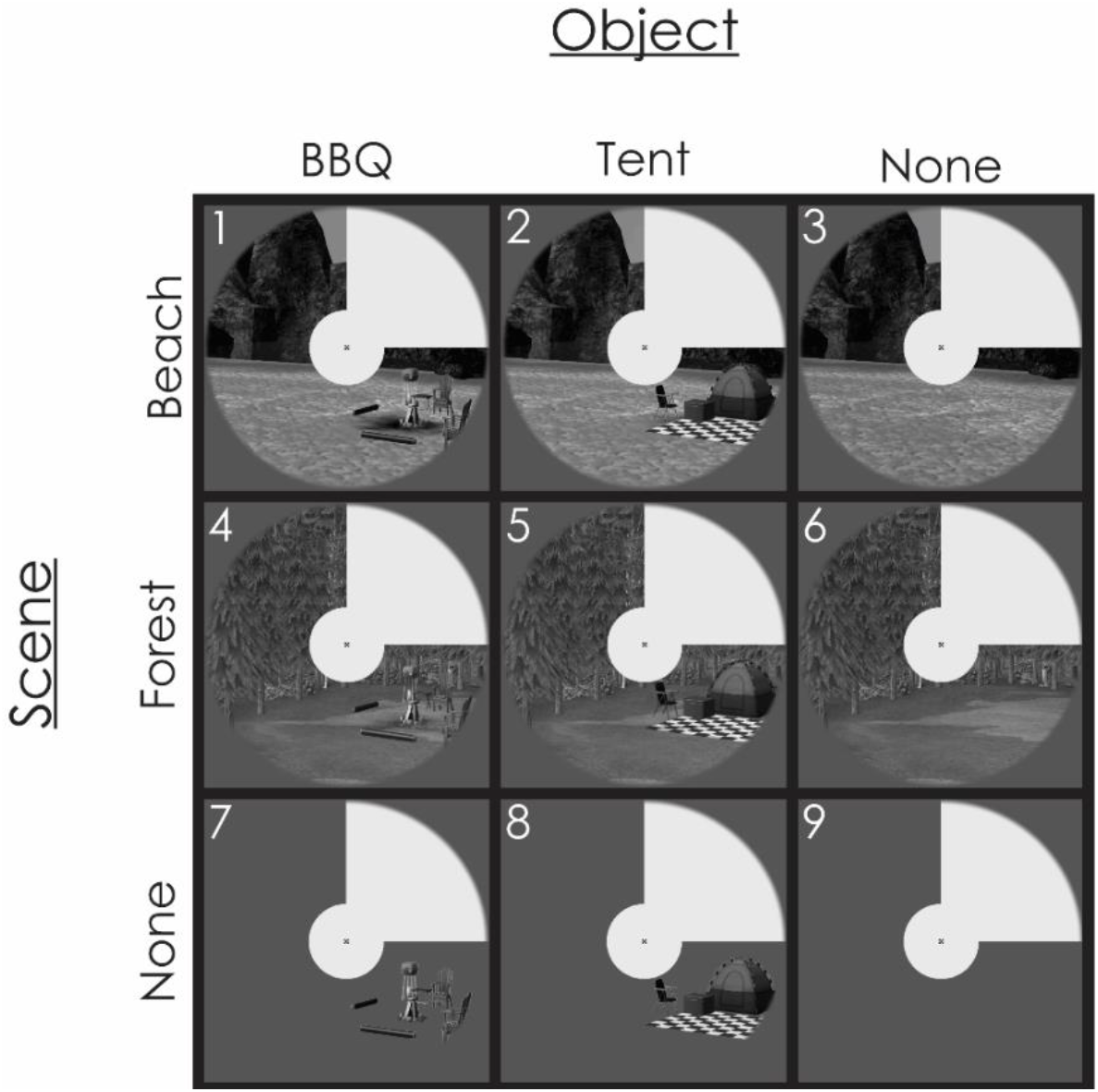
The eight stimulus images used in experiment 2. Objects (BBQ, tent) were either overlaid on scenes (forest, beach) or presented in isolation (none). Scenes either contained objects or were presented without the addition of objects (none).

In addition to classifying the object and scene identity as in experiment 1, we also classified scene backgrounds with no object (e.g., Beach) against the same scene background with an object present (e.g., Beach + BBQ). Since no fine details are required to indicate an object’s presence or absence in the image, this “object presence” feedback information may be easier to detect than the object identity – perhaps even in the periphery. We labelled our data according to scene (collapsing across objects) or according to object (identity or presence, collapsing across scenes). We performed this analysis independently for both the scene and object task data.

### Behaviour

In contrast to experiment 1, in experiment 2 participants found both tasks easy: in the scene task scoring 97.5% correct (± 1.6% stderr) and in the object task scoring 96.4% (± 2.1% stderr).

### fMRI brain imaging – MVPA

When analysing brain activity patterns from occluded fovea and peripheral regions, we were able to decode scene information regardless of task in foveal (Object Task: 54.4%, p<0.0001; Scene Task: 55.2%, p<0.0001) and peripheral V1 (Object Task: 55.7%, p<0.0001; Scene Task: 55.5%, p<0.0001, Figure 4A). Thus, reducing the object task load (as compared to experiment 1 in which participants only scored 76.6% correct), and using naturally occurring objects that the brain expects to be present in the image, might have facilitated automatic scene feedback processing widely across V1.

**Figure 4.**
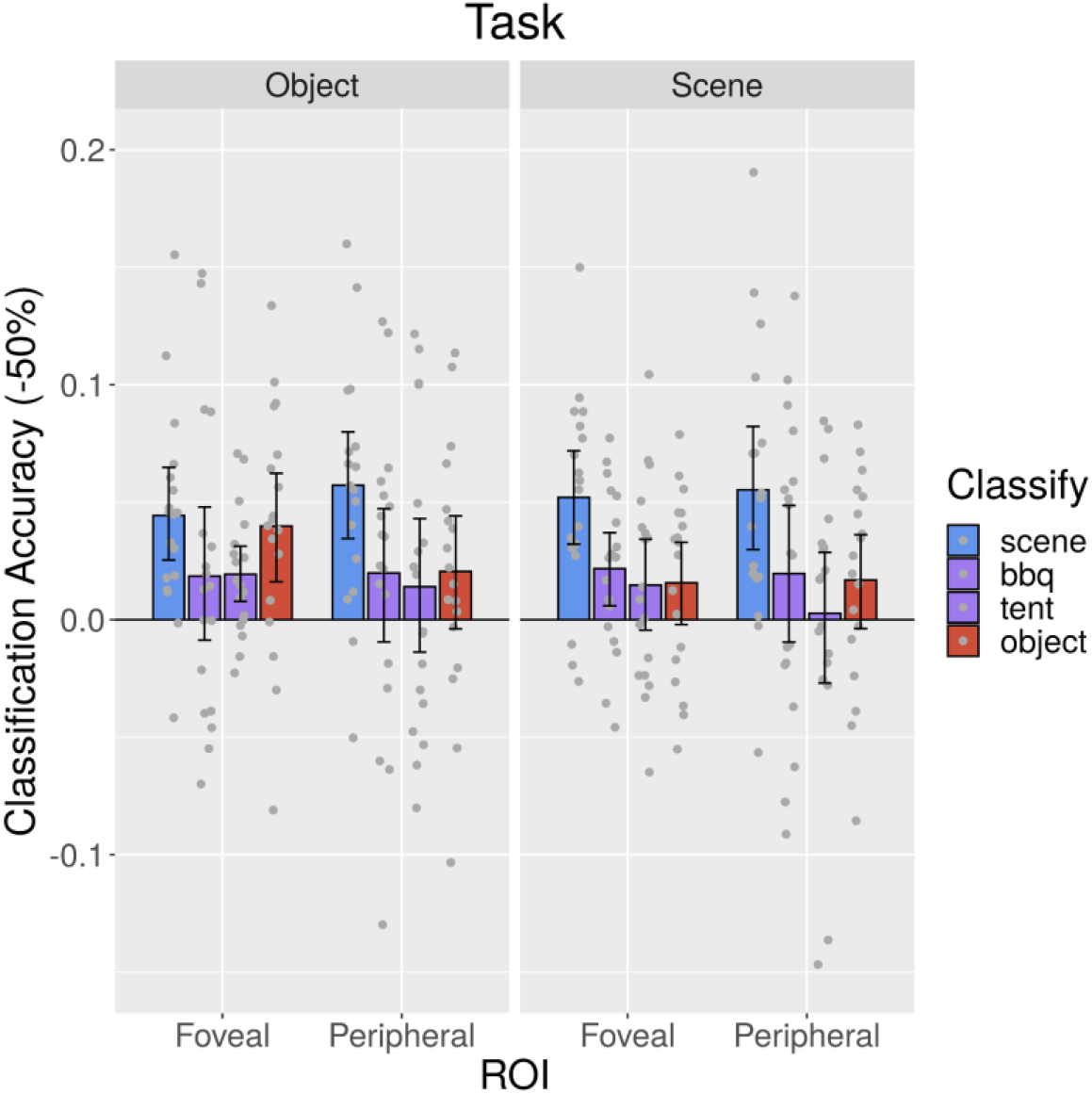
Classification accuracies for all conditions of experiment 2, in foveal and peripheral V1 regions of interest, in object and scene tasks.

We found that object presence in the scene was detectable in foveal V1 for the Tent in the object task only (BBQ during Object Task: 51.9%, p=0.0993; BBQ during Scene Task: 52.2%, p=0.0035; Tent during Object Task: 51.9%, p<0.001; Tent during Scene Task: 51.5%, p=0.0678, Figure 4B). In peripheral V1, no object information was detectable during either task (BBQ during Object Task: 52.0%, p=0.942; BBQ during Scene Task: 52.0%, p=0.0876; Tent during Object Task: 51.4%, p=0.168; Tent during Scene Task: 50.3%, p=0.858).

Object identity information was detectable in foveal V1 only during the object task (Object Task: 54.0%, p<0.001; Scene Task: 51.6%, p=0.0409). In peripheral V1 no object information was detectable during either task (Object Task: 52.1%, p=0.0409; Scene Task: 51.7%, p=0.0552, Figure 4C). This suggests that object identity is fed-back only to foveal locations in V1 when participants are asked to identify the objects in the scene. We found no significant differences in classification rates between ROIs or between different tasks.

## Discussion

In the primate brain, visual scenes and their detailed contents are processed by specialised visual cortical areas, extending from early visual cortex into ventral temporal and lateral occipital cortex (Grill-Spector and Weiner, 2014). Early and higher visual areas function together as a recurrently connected network in which cortical feedback processes contextualise and predict the feedforward information stream (e.g., Pennartz et al., 2019). To investigate the spatial organisation of cortical feedback to V1 further, we tested the pattern of feedback information to foveal and peripheral V1, during object and scene processing. We found that foveal and peripheral V1 contain high-level information that could not have arrived directly from retinal and lateral geniculate input. As such, this contextual information must have been projected to V1 from higher levels of the cortical hierarchy, or from lateral interactions within V1. We found that scene feedback information projects to peripheral V1, consistent with previous findings (Muckli and Smith, 2010; Muckli et al., 2015; Revina et al., 2018; Morgan et al., 2019; Petro et al., 2023), as well as foveal V1. This scene feedback information can be disrupted by a sufficiently difficult, orthogonal object discrimination task, but is otherwise robust and automatic when objects are predictable within the scene. We also show that the presence and/or identity of an object in the scene is fed back to foveal V1 when the visual system is engaged in an object-relevant task. This result corroborates findings from Williams (2008) of a bias for object feedback information to be projected to early foveal cortex.

Our data suggest that the spatial distribution of cortical feedback information in V1 is not a retinotopic one-to-one mapping of feedforward visual features. The pattern of scene feedback information is in line with our previous findings in which the occluded information is filled in with a “mental sketch”, similar to a simplistic line drawing (Muckli et al., 2015; Morgan et al., 2019), as well as our findings of auditory scene decoding in V1 due to cross-modal feedback (Vetter et al., 2014; Vetter et al., 2020). The pattern of feedback information for object stimuli is more complex; it is only foveal even when the objects are presented in peripheral locations, and performing a task on the objects enhances the feedback of object related information. The fact that objects are only decodable in feedback when they are relevant in terms of a task and/or synergy between objects and scene speaks for the importance of contextual prior for object perception, and in particular the role of peripheral vision to build this context. While peripheral vision is often considered as just less precise compared to the fovea, evidence is accumulating that peripheral vision has its own specific role in visual processing to fulfil (Li et al., 2021).

To explain our pattern of results for object related feedback content, we propose three potential mechanisms: feedback information related to a cognitive mental space, or active blackboard for cognitive tasks; feedback information following the retinotopic specificity and connectivity pattern of the ventral stream; feedback information related to an anticipated overt attention-shift (saccade) that would bring the to-be-focused object into the fovea.

### Cognitive active blackboard

David Mumford proposed that cortical feedback processing facilitates the function of an active blackboard, whereby expert higher areas “sketch out” what they recognise onto a lower tier visual area, similar to a blackboard, which decays over time (hence active). On such a “blackboard”, recognised features are extracted and accessible, for example the colour and the contour of an object located in the visual field. We propose that a similar mechanism could be used for cognitive tasks such as the mental comparison of two objects or for visual imagery. This account would explain previous data from Williams et al. (2008) where two objects could be compared only by using the cortical retinotopic space in early visual cortex, and such a cognitive comparison is disrupted if TMS is applied at a late time window of 350-400 ms after stimulus presentation but not earlier, consistent with a role of feedback processing after initial forward processing (Chambers et al., 2013). Our data are consistent with this idea of object feedback information, which was stronger if the task was to identify the objects, and we did not observe feedback of object information to peripheral space. In the cognitive blackboard account, cortical feedback systems support visual mental models with a spatial representation that is not restricted to the retinotopic coordinates determined during perception. Instead, V1 can be used in a flexible and adaptive way to imagine objects and scenes, using feedback signals liberated from the coordinate system in which the stimuli are presented. Another case where mental visual representations are fed back to foveal locations is Charles Bonnet Syndrome. Individuals experience complex visual hallucinations such as people, objects, or scenes because of damage to the visual system, for example, macular degeneration. These effects are similar to the notion of the early visual system as a cognitive active blackboard in the sense that V1 is involved in mental operations not necessarily triggered by feedforward inputs. Reichert et al (2013) also propose Charles Bonnet Syndrome as evidence that the cortex forms generative models of visual representations without sensory visual information.

### Cortical feedback in perception

Higher level object and scene areas are inherently biased to central and peripheral visual fields, respectively. This bias unifies the higher order and early visual cortices in an anatomically predictable scheme through retinotopic eccentricity (Hasson et al., 2003, Wang et al., 2022). Malach and colleagues propose that the functional relevance of the central-peripheral bias in higher areas is to accommodate fine detail discrimination of objects and large-scale integration of scenes, respectively (Malach, Levy, & Hasson, 2002). Higher areas have large receptive fields that make high resolution processing difficult (Lee and Yuille, 2004). Retinotopically segregated feedback might solve this problem by recruiting V1’s highest resolution capabilities (in the fovea) to scrutinise and discriminate objects (Hochstein & Ahissar, 2002). Predictive processing theories propose that higher, more specialised visual areas project their predictions of forthcoming inputs down the hierarchy to a common processing space (e.g., V1). Processing of subsequent sensory input is influenced by feedback from all levels of the hierarchy to jointly negotiate its interpretation, and potential relevance. Higher areas could use feedback processing to sketch a spatially precise interpretation of the stimulus by leveraging V1’s high spatial resolution capabilities. Such theoretical models describe the visual system as using hierarchical internal models to predict feedforward features, requiring that top-down predictions originating in higher areas restore some level of spatial precision in V1 by using its retinotopic organisation. A rough, low precision draft could, for example, indicate contour ownership by showing which side of a contour demarcates the object and which side the background. Such a code represents important contextual information that is helpful for scene segmentation, even if the precision is only a few degrees in visual angle (Petro et al., 2023). It does not help to account for the task related feedback to foveal regions in our results.

### Anticipated but suppressed eye-movements

Objects attract overt (eye-movements) and covert (suppressed eye-movements) attention shifts. Overt attention shifts result in executed saccades, which follow a process of saccade target selection determined by saliency and task relevance. Objects are salient and often selected as the location for a saccade (Kroner at al., 2019). When the task involves the comparison of two peripheral stimuli or the identification of one peripheral target object, it is conceivable that the peripheral objects are automatically selected as potential next saccade positions (Knapen et al., 2016). In our experiment, participants maintained central fixation, however saccade preparation might have already initiated the process of saccadic remapping. This process would project the object that would have been the subject of a saccade into the anticipated post-saccadic cortical location, in this case, the fovea. We have shown previously that cortical predictive mechanisms are fast enough to project internally generated predictions to the anticipated retinotopic location of post-saccadic input (in the case of a motion illusion, Edwards et al., 2017). In the current data, the decoding of object information from foveal locations could be related to an anticipated retinotopic remapping.

Our three accounts are not mutually exclusive. With regards to scene processing, predictive computations during perception of natural images might be automatic. Here, our prior knowledge about scenes allows us to perceive the gist whilst explaining away the finer details that do not require attention. During the object task, higher-level object-selective areas might project their hypotheses about object features to early visual foveal areas where higher-resolution capabilities can assist in resolving ambiguities. Here, additional cognitive processes such as attention recruit early visual areas, potentially to enhance the neuronal response to stimuli that might be the target of an upcoming saccade.

## Conclusion

We present the feedback information processing in V1 during the disambiguation and recognition of objects and scenes, possibly from functionally specialised higher areas. We propose that the role of V1 should be thought of as a functionally flexible extension of higher areas, whereby V1’s high-resolution processing might be recruited adaptively according to task requirements. Our finding opens several intriguing avenues of research. If object-related feedback targets the fovea, does this imply that the visual imagery of objects activates foveal V1 specifically? If so, would it mean that imagery that is spatially constrained to periphery (e.g., picturing a face in the left periphery) is more challenging? Another straightforward prediction from foveal object feedback is a bias under which an object presented in the periphery should be remembered as more central (a framing effect like boundary extension, which makes objects smaller and more foveal). Finally, if we follow the hypothesis that higher areas are tuned to object invariance e.g., rotation/translation, feedback signals from these areas should then also be “generic”, which would result in the prediction that feedback is biased not only towards foveal presentation, but also canonical object presentation or prototypical exemplars of object categories.

## Abbreviations

fMRI: functional magnetic resonance imaging
MVPA: multi-variate pattern analysis, ROI, region of interest
pRF: population receptive field
V1: human primary visual cortex

## Data and availability

The data and code are available here: https://github.com/Matt-A-Bennett/scene_objects_analysis.

## CRediT authorship contribution statement

**Matthew Bennett:** co-design, data acquisition, data analysis, figure preparation, writing. **Lucy Petro:** co-design, co-supervision, writing. **Clement Abbatecola:** writing. **Lars Muckli:** design, supervision, funding acquisition, writing.

## Acknowledgements

This project has received funding from the European Union’s Horizon 2020 Framework Programme for Research and Innovation under the Specific Grant Agreement No. 720270, 785907, and 945539 (Human Brain Project SGA1 SGA2, and SGA3), and European Research Council (ERC StG 2012_311751-’Brain reading of contextual feedback and predictions’, both awarded to LM, and Biotechnology and Biological Sciences Research Council (BBSRC BBN010956/1) ‘Layer-specific cortical feedback’ to LM (with LSP). We thank Frances Crabbe for assistance with data collection, and Angus Paton for comments. For the purpose of open access, the author(s) has applied a Creative Commons Attribution (CC BY) licence to any Author Accepted Manuscript version arising from this submission.

## Methods

### Experiment 1

#### Participants

We recruited ten healthy participants (4 male; mean 22.9 years old, range = 18-29) using the University of Glasgow, School of Psychology subject pool. All participants had normal or corrected eyesight and no history of brain damage. Participants gave consent and were screened in accordance with the safety criteria for fMRI scanning. The study was conducted in accordance with approval from a local ethics committee (#CSE01326).

#### Stimuli and Apparatus

We projected the stimuli with a refresh rate 60 Hz at a resolution of 1024 × 768 onto a screen (19.0° × 14.2° visual angle) which participants viewed via a mirror attached to the head coil. We programmed the experiment and displayed it using Presentation (Version 16.5). There were four experimental conditions and five retinotopic mapping conditions (‘periphery’, ‘periphery border’, ‘fovea’, ‘fovea border’, and object locations, Figure 1B). In addition, participants underwent polar angle and eccentricity retinotopic mapping to localise V1. The images were one of two grayscale natural scenes (‘Mountain’ and ‘Seaweed’) with a pair of superimposed abstract objects belonging to one of two object categories (‘Cubic’ and ‘Smooth’, Figure 1). Each object appeared at 6.1° eccentricity. The width/height of the objects was approximately 2.0°. The objects were centred atop a 3.5° diameter black disk. The objects were created using a custom algorithm written in Matlab. Specifically, each object shown throughout the experiment was unique (i.e., no object appeared more than once: 1728 unique objects were created in total) and generated by manipulating 4 main feature dimensions (mainly related to the number, size, shape, and rotational position of the objects’ protrusions). Thus, objects could not be discriminated by looking at only a small part of each object, and more than one location of the object had to be considered to attain good discrimination performance. We aimed to make them similar to those used in Williams et al. (2008). Each image spanned 19° x 14.2°. We occluded central (3.8° x 3.8°) and lower right (9.5° x 7.1°) image portions. To control for low level image properties in those areas of the scenes visible to participants, the SHINE MATLAB toolbox (Willenbockel et al., 2010) was used to match the luminance histograms (i.e. scale the pixel intensity such that there are same number of pixels at each intensity for each scene – this also matches the average/global luminance) and the amplitude spectrum of each image was replaced with the average spectrum of the two images, thus exactly matching the images at each spatial frequency and orientation.

#### Task and Procedure

Participants completed 12 functional runs across two days (6 runs per day). In odd or even runs (counterbalanced across subjects), subjects performed one of two tasks: 1) judging whether the two objects were identical or not (50% identical) or 2) indicating the scene identity when cued by a frame that was slightly blurred (8 blurred frames per run). Subjects used the index and middle finger of the right hand, and mapping was counterbalanced across subjects. Subjects maintained fixation throughout the experiment. Each run consisted of 12 blocks. Each block consisted of 8 trials drawn from the same condition (e.g., Mountain + Cubic objects). The block condition order was pseudo-randomized for each subject, with the constraint that no condition was repeated twice in a row. Each trial consisted of a stimulus flashed on and off (4 Hz) for 1s followed by a 1s fixation period during which participants responded, thus each block lasted 16s. A 12s baseline period preceded each stimulation block. At the end of the baseline period following the last stimulation block, the retinotopic mapping period of the run started. The retinotopic mapping period consisted of 10 blocks, each lasting 12s, of contrast reversing checkerboard stimuli (4 Hz) at one of five locations (central, central-surround, periphery, periphery-surround, object locations) - there were two blocks per location. A 12s baseline period preceded each checkerboard stimulation block (and also immediately followed the last block). Each run lasted 9 mins 55s. In addition to the task runs described above, each scanning session ended with either polar angle or eccentricity retinotopic mapping. Each scanning session lasted approximately 1.5 hours.

#### Data Acquisition

We acquired functional and anatomical MRI data using a 3 Tesla MRI system (Siemens Tim Trio) with a 32-channel head coil. For the functional scans an echo-planar imaging sequence was used with the following parameters: 18 slices, aligned with the calcarine sulcus, gap thickness 0.3mm, TR-1s, TE-30ms, 588 volumes per run (polar angle mapping = 808 volumes, eccentricity mapping = 553 volumes), a FOV of 220mm, flip angle of 62° and a resolution of 3.0mm^3^. The anatomical MRI sequence had a TR of 2.3s, 192 volumes, and a resolution of 1.0mm^3^.

#### Data preprocessing

The functional and anatomical data were pre-processed using BrainVoyager QX 2.8 (Brain Innovation). We discarded the first two volumes of each functional run to avoid saturation effects. We corrected the functional data for each run for slice acquisition time and head movements. We removed linear and low frequency drifts in the data. We then aligned the functional data with the high-resolution anatomical data and transformed into ACPC space. We created a model cortical surface of the white matter grey matter boundary from the ACPC anatomical scans.

#### ROI Definitions

For each subject, we projected the functional data onto the cortical surface. After localising V1, using the polar and eccentricity data, we defined the ROIs by contrasting the appropriate target mapping stimuli with the corresponding surround mapping stimuli (see fig 1B, and 1C). Custom Matlab algorithms were used to estimate the population receptive fields (pRF, see Dumoulin & Wandell, 2008) from the polar angle and eccentricity data for all V1 and early visual foveal voxels, allowing us to reject from the occluded ROIs voxels with population receptive fields (pRFs) falling outside the occluded regions (see fig 1A, lower right).

#### Data Analysis

A GLM was used to estimate each voxel’s HRF response for each block. We attempted to classify either the scene or object identity using a linear Support Vector Machine (SVM) classifier. This analysis was performed independently for the scene and objects task runs. Specifically, we trained an SVM to classify the Scene or Object using the associated multivariate voxel response patterns. For example, to classify the Scene we labelled the 4 conditions according to the background scene so as to create a binary classification problem (i.e., Mountain + Cubic, Mountain + Smooth labelled ‘1’ vs. Seaweed + Cubic, Seaweed + Smooth labelled ‘2’). A corresponding approach was taken when classifying Object Identity. We used a ‘leave one run out’ cross-validation procedure.

Since the sensitivity of the SVM varies with the number of features entered, it was important to precisely control for the different number of voxels in the ROIs (foveal: 98.3±41.1 std dev, peripheral: 49.0±16.7 std dev). For each ROI we therefore took equally sized random samples of voxels and ran the SVM. The number of voxels sampled was set per subject as 75% of the number in the peripheral ROI (since this was always the smallest ROI). We did this 1000 times and averaged the SVM results to get a single accuracy value. This sampling approach has the added advantage that our SVM accuracy estimates are less susceptible to being skewed by any particular combination of voxels.

To assess the significance of group level effects, the mean group classification value was bootstrapped 100’000 times, and a p-value was calculated as the proportion of bootstrapped classification values falling above the observed value. In each experiment, we applied a Bonferroni correction to account for multiple comparisons (the alpha in experiment one was 0.05/8=0.006; in experiment two it was 0.05/16=0.003).

### Experiment 2

Unless stated, the details of experiment two were identical to experiment 1.

#### Participants

We recruited eighteen healthy participants using the University of Glasgow, School of Psychology subject pool. All participants had normal or corrected vision and no history of brain damage. We screened participants for MRI safety and acquired consent. The study was conducted in accordance with approval from a local ethics committee (#CSE01326).

#### Stimuli and Apparatus

The stimuli were projected at a resolution of 768 × 768 onto a screen (19.9° × 19.9° visual angle). There were eight experimental conditions and four retinotopic mapping conditions. Our images (see figure 3) were created by combining one of three background possibilities (Beach, Forest, None) with one of three object possibilities (BBQ, Tent, None). Thus, there were nine possible image combinations. This resulted in eight experimental images and one blank image used as a fixation condition (None/None). Each image was circular, with a radius of 9.9°. All nine images were occluded in a similar way to experiment 1 (central radius 2.6° and upper right quadrant). The mean luminance of every image was the same (85). In particular, in each image the mean luminance of the set of pixels belonging to the background scene and object were also the same when considered separately (85).

#### Task and Procedure

Participants completed 10 functional runs across two days (5 runs per day). Participants performed either an object or scene identification task on alternate days (counterbalanced across participants). The object task was to identify the object in the image (BBQ, Tent, or None). Similarly, in the scene task participants had to identify the background scene in the image (Beach, Forest or None). Participants made button presses with the index, middle or third finger of the right hand (mapping counterbalanced across runs each day for each subject). Participants responded at any point during the 12 second block. Each run consisted of 24 blocks. Blocks consisted of a stimulus flashed on and off (4 Hz). Participants responded any time after the onset of a block. The occluder did not flash and remained present throughout the entire run. The retinotopic mapping period consisted of 4 blocks, each mapping one of four locations (central, central-surround, periphery, and periphery-surround). Each run lasted 11 mins 24s.

#### Data Acquisition

For the functional scans an echo-planar imaging sequence was used with the following parameters: 31 slices, aligned with the calcarine sulcus, gap thickness 0.2mm, TR-2s, TE-30ms, 342 volumes per run (polar angle mapping = 396 volumes, eccentricity mapping = 268 volumes), and a resolution of 2.0mm^3^.

#### ROI Definitions

Since we did not have reverse-checkerboard mapping stimuli for the object locations, V1-object ROIs were defined by contrasting object only experimental conditions with all the other conditions.

#### Data Analysis

A Support Vector Machine (SVM) was used to classify the Scene, Object Presence or Object Identity in the VI ROIs. For example, to classify the Scene we labelled the 6 conditions that contained a background scene so as to create a binary classification problem (i.e., Beach, Beach + BBQ, Beach + Tent labelled ‘1’ vs. Forest, Forest + BBQ, Forest + Tent labelled ‘2’). A corresponding approach was taken when classifying Object Identity. In the case of classifying Object Presence, we labelled the two object-free scenes as ‘1’ and then classified them against the same two scenes with an embedded object. We did this independently for the presence of BBQ and Tent in the scene. The number of voxels in the ROIs were V1-Object: 228.0±27.2 std dev, foveal: 475.2±117.0 std dev and peripheral: 116.6±41.1 std dev. These disparities were dealt with as for experiment 1.

